## Supplementary material for "RNA structure prediction using positive and negative evolutionary information": README to expand supplemental materials

Instructions to open the supplemental materials from the bioRxiv
------------------------------------------------------------------

(1) download Supplementary Material file from
     https://doi.org/10.1101/2020.02.04.933952

(2)  A file named 'media-1.gz' will download to your computer

(3) To open the tarball, in your terminal use the command:

    tar xf media-1.gz

    that will expand the directory
    
    supplemental_material/


(4) Methods that DO NOT work:

    (a) unzipping the file using the GUI (graphical user interface) 
    (b) gunzip media-1.gz
 
