## supplemental_material for "RNA structure prediction using positive and negative evolutionary information": CHANGES.txt

- 224/ARRPOF-assoc.sto : slight improvements to alignment.
- 104/hopC.sto : alignment withdrawn since an alternate alignment with better sequence conservation would invalidate the evidence of covariation.
- 224/DUF1646.sto : improved P2 in a few sequences where it previously fit poorly
- 224/nifS.sto : alignment withdrawn due to a total overlap with the much better skipping-rope motif
- 22/anti_hemB.sto : touched up alignment in middle stem.
