## supplemental_material for "RNA structure prediction using positive and negative evolutionary information": README.txt

This is a repository of multiple-sequence alignments from work by Zasha Weinberg.  Most of these alignments have been submitted to the Rfam database.

** STATUS OF ALIGNMENTS **

Most alignments were submitted to the Rfam database.  However, for some alignments, it was considered ambiguous whether the alignment was likely to represent a biologically function RNA (or ssDNA) molecule or not.  These alignments were not submitted to Rfam.  The file 'not-for-Rfam' lists these alignments.  Not all alignments that were submitted to Rfam were incorporated into this database.

In some cases, it was not possible to recover the exact alignment submitted to Rfam, or associated with a given paper.  In this case, a comparable alignment was substituted.

In some cases, a change to the alignment was intentionally made after publication to improve the alignment, and such alignments are indicated with a #=GF STATUS tag in the file, which explains the changes.  These changes are summarized in the file CHANGES.txt

** FILES WITH SUMMARY OR META DATA **

The tab-delimited file 'PAPERS' gives a link to the paper corresponding to each subdirectory.

The file 'MISSING' is notes on alignments that are not in this repository, but could be.

** DIRECTORY STRUCTURE **

Each subdirectory contains alignments from a different paper.  However, the directory 'patches' corresponds to miscellaneous alignments of known RNAs that were used to find homologs that are (or were) not detected by existing Rfam alignments.  These alignments are rough and generally contain only a part of the RNA -- their purpose is simply to be able to annotate more of the already known RNAs.

** FORMAT OF ALIGNMENTS **

All alignments are in Stockholm format, and predominantly use conventions established by Rfam.

Pseudoknots: Pseudoknots are represented by matching upper- and lower-case letters in the #=GC SS_cons line, as in the Rfam database.  

Sequence names: As in Rfam, the names of each sequence in the alignments is of the form SEQID/START-END, where SEQID is a sequence accession, START is the coordinate of the 5'-most nucleotide of the aligned sequence and END is the 3'-most nucleotide.  The file SAH/SAH-from-Wang-etal.sto is, however, in a different format.  The sequence accessions refer to RefSeq or environmental metagenomic sequences from various sources.

"Key" nucleotides: alignments in the 'variants' subdirectory (see https://www.ncbi.nlm.nih.gov/pubmed/28265071) have additional information defining the "key" (presumed ligand-binding) nucleotides that are varied.  A
#=GC CORE
line defines the "key" nucleotides (i.e., nucleotides that are presumably in the ligand-binding core).  Columns with the key nucleotides are indicated with an 'X' in this line.
A 
#=GF CORESEQ
tag defines the identities of the nucleotides for the given variant.  The nucleotides in #=GF CORESEQ correspond, from 5' to 3', to the columns marked with 'X' in the #=GC CORE line.  Homology searches for these variant alignments could restrict the predicted homologs to those having the nucleotides in #=GF CORESEQ in the columns indicated by #=GC CORE.

** OTHER FILES **

The following files were not mentioned above.

The file 'commands.txt' is a note on commands to run to generate automated data, for my convenience.

The tab-delimited file 'sto-seqid-taxid.tab' associated sequence IDs used by alignments within the repository with NCBI taxonomy IDs, where possible for sequences from the RefSeq nucleotide database.  The fields are: (1) the sequence accession referring to the RefSeq or other nucleotide database, and (2) the taxonomy ID (or -1 if the sequence accession is not in Refseq).  The alignment SAH/SAH-from-Wang-etal.sto is not reflected in this file.

README.txt : this file
